# Extraintestinal survival and host immune response to *Vibrio cholerae*

**DOI:** 10.1101/2022.11.03.515087

**Authors:** Foster K. Agyei, Nana E. Adade, Nsoh G. Anabire, Vincent Appiah, Yaw Bediako, Samuel Duodu

**Author notes:** Corresponding author; (SD).

## Abstract

*Vibrio cholerae* is best known to cause the deadly disease cholera. However, in recent years this bacterial pathogen has been found to invade intestinal layers and translocate into the bloodstream of humans. The aim of this study was to investigate the molecular basis of *V. cholerae* bacteremia. Nine (9) strains of *V. cholerae*; six (6) environmental strains of non-O1/non-O139 serogroup and three (3) clinical strains of O1 serogroup and El-Tor serotype were screened for survival in serum obtained from immunocompromised patients. Serum from immunocompetent individuals with no known underlying conditions were used as healthy controls. Five (5) environmental strains and one (1) clinical strain of *V. cholerae* were identified to survive the bactericidal action of serum. Whole genome sequence analysis revealed the cholix toxin (*ChxA*) and genes encoding for siderophores *(FepE* and *EntD*) as possible virulence factors used by the environmental strains to cause invasive bloodstream infection. Peripheral blood mononuclear cells (PBMCs) stimulated with *V. cholerae* revealed increased expression of some cytokines; IL-1ß and IL-13 and the chemokine; RANTES especially among diabetics. The present study illustrates the potential survival of *V. cholerae* in blood, which could be aided by scavenging for iron from their host leading to severe infections.

## 1. Introduction

Pathogenic *Vibrio cholerae* strains are the causative agents of cholera, a serious diarrheal disease with a high mortality rate [1]. The strains of *V. cholerae* are classified into two biotypes (El Tor and classical) and over 200 serogroups [2]. The biochemical properties and phage sensitivity of the El Tor and classical biotypes are used to identify them while serogroups are distinguished by the structure of their O-antigen. *V. cholerae* serogroups O1 and O139 are toxigenic, expressing virulence factors found to be essential for pandemic spread. Among these are the cholera toxin (CT), a potent enterotoxin responsible for the profuse rice water diarrhoea typical of the disease and the toxin co-regulated type IV pilus (TCP), which is important for intestinal colonization [3]. The non-toxigenic *V. cholerae* strains (non-O1/non-O139), lacking the *ctx* gene that encodes for CT production have only been linked to a few cases of cholera, but known to cause bacteremia and other extraintestinal infections particularly in immunocompromised people [4-7].

To survive in a host, an invading bacteria must be able to detect and rapidly adapt to changes in growth conditions between the external environment and the host, as well as inside the host’s many microenvironments. Bacterial pathogens generally produce virulence factors, such as adhesins, capsules, iron uptake systems and toxins, to evade host responses and to multiply and spread within the host. Both CT and TCP have not been associated with host invasion. However, the presence of type 3 and type 6 secretion systems in some *V. cholerae* strains [8, 9], enables effector proteins or toxins to be translocated directly into the host cytosol. Other cytotoxic proteins with potential invasive properties including haemagglutins, haemolysin, TagA proteases and phospholipases have been suggested to play significant roles in cholera pathogenesis particularly in environmental non-toxigenic strains.

Presently, how the non-O1/O139 cause infection, alongside their survival in serum remain unclear [10]. The role of the immune system in such infections is also important as the rapid recognition of the invading bacteria by immune cells plays a key role in preventing the pathogen from harming the host. Although it has been shown that naturally existing IgG recognizing *V. cholerae* outer membrane protein U (OmpU) exerts a serum-killing activity in a complement C1q-dependent manner [10], the immunological responses induced during *V. cholerae* bacteremia are not well characterized. To address these gaps, we investigated for the presence of genes that support *V. cholerae* survival in blood as well as characterized the specific immune responses elicited by the host to fight this infection.

## 2. Materials and Methods

### 2.1. Collection of bacterial isolates

Six (6) environmental and three (3) outbreak *V. cholerae* isolates were used in this study. Environmental isolates were isolated from environmental water samples. To recover *V. cholerae* from the environment, water samples were collected aseptically into sterile bottles containing 5mL of alkaline peptone water (pH 8.5) at different times from different points of the Korle Lagoon, Accra, Ghana from 2016 to 2020. These were cultured on Thiosulphate Citrate Bile-salt Sucrose (TCBS). Archived outbreak strains selected from four different outbreak years (2010, 2011, 2014, 2015) were retrieved from storage at -80°C from the Public Health Reference Laboratory (NHPRL) in the Greater Accra region of Ghana and revived by thawing at 37°C before inoculating on TCBS. To confirm the identity of the isolates a loop full of enrichment culture was collected and streaked onto TCBS agar plates and incubated at 37°C for 18-24h [11]. Presumptive colonies were sub-cultured onto Mueller Hinton agar plates for 24 hrs to obtain pure cultures. DNA was extracted using bacterial Zymogen DNA extraction mini kit from the isolates and subjected to conventional PCR amplification for the detection of the outer membrane protein gene (*ompW*) confirming them as *V. cholerae*. Toxigenicity was further tested by PCR targeting the cholera enterotoxin subunit A gene (*ctxA*) [12]. Isolates with non-agglutinating abilities and lacking the virulence genes encoding for CT in toxigenic strains were considered as non-O1/non-O139 strains [13]. Oligonucleotide primers (Table 1) were synthesized by Inqaba Biotechnical Industries, South Africa.

**Table 1.**
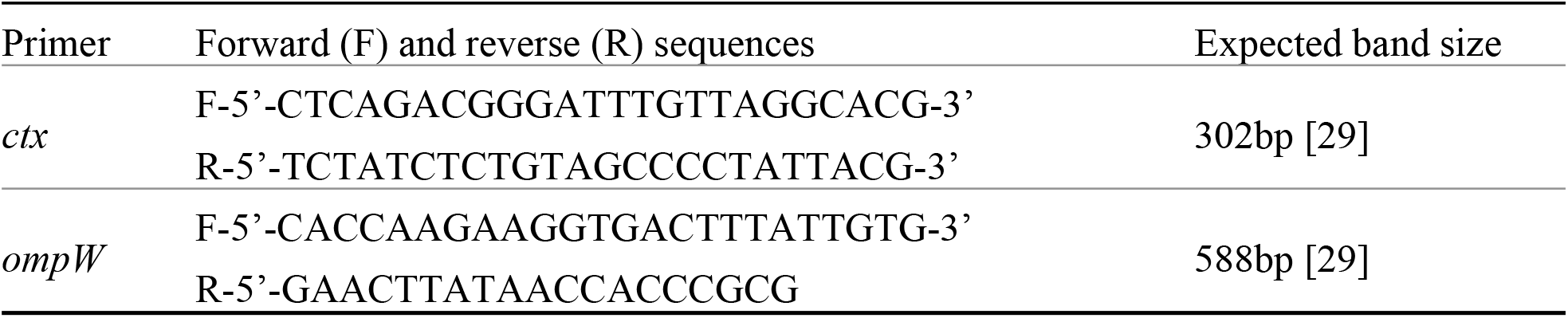
Primer list for PCR Amplifications.

### 2.2. Blood sample collection and separation of human serum

A volume of 8mL blood samples from study participants were collected from Diabetic and Liver diseases clinic at Lekma Hospital. Venous blood samples from patients were taken with the help of a laboratory scientist. Samples were collected into BD vacutainer tubes containing sodium heparin. The closed tube was inverted carefully several times to ensure proper mixing with anticoagulants. Samples were transported and stored at a temperature of 2 ^º^C-10 ^º^C for a maximum of 5 days. Whole blood samples were collected into red-top tubes with no anticoagulants. Blood was allowed to clot and centrifuged at 4000rpm for 10 minutes. Serum was collected using a sterile pipette and transferred into a sterile Eppendorf tube. Before further processing of samples, whole blood genomic DNA were screened for the presence of *V. cholerae* and other bacterial pathogens by PCR targeting the *ctx* [14], *ompW* [14] and *16S rRNA* [15] genes.

### 2.3 Determination of *V. cholerae* survival in serum

Experiments to determine pathogen survival were performed in human serum. The assays were performed using both clinical and environmental isolates of *V. cholerae*. In all, nine (9) strains of *V. cholerae*; six (6) environmental strains of non-O1/O139 serogroups and three (3) clinical strains of O1 serogroups of El Tor (C22, C17) and classical (C6) Ogawa biotypes were screened for survival in serum obtained from 50 study participants. For each bacterial strain, three colonies of pure culture were transferred into 5mL LB broth in a 15mL falcon tube and grown at 37^º^C with shaking overnight. A volume of 150μl of the overnight culture was then transferred in 5mL fresh LB medium and grown to an optical density of 0.5 (OD_600_). Bacterial culture was diluted to 1 in 1000 in Phosphate Buffered Saline (PBS) and 50μL of the culture was plated to determine the number of viable cells prior to serum exposure. About 50μL of bacteria culture was added to 100μL of serum in a sterile Eppendorf tube and incubated for 4 hours at 37^º^C. Following incubation, 50μL of culture was plated on MHA/LB agar. Growth in serum was determined by calculating the differences in bacterial CFU without serum (T_0_) and after 4 hours (T_4_) incubation of bacterial/serum mixture. Percent survival was calculated using the formula; [(T_4_/T_0_) ×100]. Experiment was performed in three technical replicates [16].

### 2.4 Heat Inactivation of serum and *V. cholerae* survival

The protocol for serum heat inactivation was adopted from the work by Fante et al. [17]. Briefly, frozen serum samples were thawed at room temperature and subsequently heat inactivated in a pre-heated water bath at 56 °C for 30 minutes. Bacterial growth in native and inactivated serum was evaluated using the same protocol as described above to assess the impact of complement activity on *V. cholerae* survival.

### 2.5. Genomic DNA extraction

DNA of surviving strains were extracted using Zymo Fungal/Bacterial and QIAamp DNA miniprep Kits following the manufacturer’s protocol. Briefly, about 10^8^ bacterial cells was resuspended in 200μl of PBS and added to a ZR BashingBead Lysis Tube. A volume of 750μl BashingBead Buffer was added to the tube. The tube was placed in a bead beater with a 2ml tube holder assembly and homogenized for 20 minutes at maximum speed at 10,000×g for 1 minute, and 400μl of supernatant was transferred to a Zymo-Spin filter in a collection tube. This was centrifuged at 8000×g for 1 minute and the filter was washed with 200μl of DNA pre-wash buffer followed by additional step with 500μl g-DNA Wash Buffer. The Zymo-Spin column was then transferred to a clean 1.5ml microcentrifuge tube and genomic DNA was eluted using 100μl DNA elution buffer. The ultra-pure DNA was quantified using Nanodrop and kept at -20 °C for sequencing later.

### 2.6. Whole genome amplification and sequencing of *V. cholerae* strains

Whole genome sequencing was performed using next-generation-sequencing on Illumina Hiseq analyser platform. Genome sequencing was done by MicrobesNG (Birmingham) using the Nextera XT library prep kit (Illumina) for genomic DNA library preparation, following the manufacturer’s protocol with the following modifications: two (2) nanograms of DNA were utilized as input instead of one (1) ng, and the PCR elongation duration was extended from 30 seconds to 1 minute. A Hamilton Microlab STAR automated liquid handling system was used for DNA measurement and library preparation. On a Roche light cycler 96 qPCR machine, pooled libraries were quantified using the Kapa Biosystems Library Quantification Kit for Illumina. A 250bp paired end technique was used to sequence the libraries on the Illumina HiSeq. Reads were adapter trimmed using sickle 1.33 with default parameters [18] *De novo* assembly was performed on samples using SPAdes version 3.14 [19] and Contigs were annotated using Prokka 1.11 [20] all via MicrobesNG (Birmingham).

### 2.7. Analysis of whole genome sequencing data

The genomes of the strains were further annotated with the help of the Rapid Annotation Subsystem Technology (RAST) program [21]. Nucleic acid Fasta files from the annotation results were used to perform a pangenome analysis for the presence and absence of genes among the *V. cholerae* strains with the help of the Prokka and Roary software. Sequence data for the individual isolates used in this study is deposited at NCBI (Sequence Read Archive) with accession numbers ranging from SAMN26804320-31.

### 2.8. PBMC isolation and thawing

Four millilitres (4mL) of whole blood were diluted with equal amounts of HBSS without Ca^+^, Mg+ and Phenol red in a 1:1 ratio and thoroughly mixed. Diluted blood was gently laid on top of four millilitres (4mL) Ficoll-paque premium in a 15mL conical centrifuge tube and spun for 30 minutes at 500×g at room temperature. PBMCs were gently aspirated using a sterile Pasteur pipette and transferred into a fresh tube. Isolated PBMCs were washed twice with RPMI by centrifugation for 10 minutes at 800rpm. The PBMCs were resuspended in 500μl of RMPI and counted using an automated cell counter. A freezing medium (90% FBS + 10% DMSO) was added to cells, transferred to cryovial tubes and stored at -80°C freezer overnight before transferring to liquid nitrogen tank. Cryovials containing PBMCs were removed from the liquid nitrogen tank, equilibrated on ice and thawed by immersing in 37°C water-bath with gentle swirling for 2 minutes. The exterior of cryovials were wiped with 70% ethanol to limit the possibility of contamination. The PBMCs were gradually transferred to a 15ml centrifuge tube in order to avoid osmotic shock prior to addition of 10ml of pre-warmed cell culture medium (RPMI). The cells were then centrifuged to remove the freezing medium containing DMSO and the supernatant discarded. The cells were then resuspended in RPMI and their viability and yield determined using an automated cell counter (Table 2).

**Table 2.** List of Donor and Their Respective PBMC Yield

| Donor | Concentration of Cells/ml |
| --- | --- |
| Donor 1* | $1.1 \times 10^6$ |
| Donor 2* | $4.4 \times 10^5$ |
| Donor 3** | $3.4 \times 10^6$ |
| Donor 4** | $2.1 \times 10^5$ |
| Donor 5*** | $6.1 \times 10^5$ |

\*Diabetic patients \*\*Liver disease patients \*\*\*Healthy control

### 2.9. Cell stimulation

In the cell stimulation assay, bacteria cells and PBMCs were resuspended in RPMI. The negative control wells contained cells with distilled water only whilst the positive control wells consisted of PBMCs with Phytohemagglutinin (PHA). The ratio of PBMCs to *V. cholerae* cells used for the stimulation was 50,000:100,000 (1:2). The Bacterial culture wells were resuspended in RPMI and added to a 96 well microtiter plate containing PBMCs. The negative control wells were PBMCs only with bacteria, and cells with PHA were used for positive control.

### 2.10. Detection of immune cell components using human cytokine magnetic 25-plex panel

For the cytokine multiplex assay, 2 standards comprising the human 16-plex and human 14-plex standard were used. To detect analytes (IL-10, IL-12, IL-13, IL-15, IL-17A, IL-1 beta, IL-IRA, IL-2, IL-2R, IL-4, IL-5, IL-6, IL-7, IL-8, IP-10, MCP-1, MIG, MIP-1 beta, MIP-1 alpha, RANTES, TNF alpha), 1× biotinylated detector antibody was prepared in a conical tube. For a single assay well, 100μl of biotin diluent and 10μL of 10× biotinylated antibody was added. The liquid was decanted, and the wells were washed twice with 200μL 1× Wash solution. A volume of 100μL of 1× biotinylated detector antibody was added to all assay wells and the plate was covered and incubated for 1 hour at room temperature on an orbital plate shaker. 1× Streptavidin-RPE solution was prepared and 100μl was added to each assay well. The plate was covered and incubated for 30 minutes at room temperature on an orbital shaker. The liquid was decanted, and wells were washed 3 times with 200μl 1× Wash solution. In reading the assay results, 150μl of 1× Wash solution was added to each assay and placed on an orbital plate shaker for 2-3 minutes prior to analysis. The plate was uncovered and inserted into the XY platform of the MAGPIX instrument to determine the concentrations (pg/mL) of the samples.

### 2.11. Statistical analysis of data

An analysis of variance (ANOVA) test was performed on the data using the R software. The test was done at a 0.05 level of significance. P < 0.05 was statistically significant.

## 3. Results

### 3.1. Study participants

A total of 50 patients including healthy controls consented and were recruited into the study between February and June 2021 (Table 3). Seven participants suffering from liver disorders that tested positive for Hepatitis B viral infections were on antiviral therapy and the 3 participants with liver diseases because of lifestyle choices were on lifestyle modification therapy. For those suffering from diabetes, all participants had been subjected to insulin injection and dietary management.

**Table 3.** Demographics and Clinical History of Study Participants

| Disease | Liver Disorders | Diabetes | Healthy Controls |
| --- | --- | --- | --- |
| Gender | Male = 8 | Male = 10 | Male = 5 |
|  | Female = 2 | Female = 20 | Female = 5 |
| Age | Male = 20 – 47 (years) | Male = 29 – 65 (years) | Male = 22 – 27 (years) |
|  | Female = 18 -55 (years) | Female = 34 –71 (years) | Female = 23 – 40 (years) |
| Therapy | Antiviral medications,<br>lifestyle modification | Dietary management,<br>insulin injection |  |

### 3.2. Environmental isolates of *V. cholerae* survive better in human serum compared to clinical isolates

In the first step of the study, we tested for the resistance of *V. cholerae* to human serum. Human serum is thought to be a crucial host defense against bacterial infections because of its principal component, the complement. As a result, invasive bacteria’s resistance to serum killing is recognized as an essential pathogenicity feature. Human sera from 50 individuals comprising; diabetic, liver disease and healthy individuals (confirmed to be negative for presence of bacteria by PCR) were screened with environmental non-O1/O139 *V. cholerae* isolates in comparison with clinical O1 strains. Clinical strains of *V. cholerae* generally are not considered to be a cause of *V. cholerae* bacteremia but few reports have implicated these strains as rare causes of *V. cholerae* bacteremia [22]. Tables 4 and 5 show the survival and percentage survival rates, respectively of all the isolates after serum exposure. Environmental isolates recorded highest survival rates compared to clinical strains. Among the environmental isolates, E4 survives better and was able to withstand serum bactericidal action from at least one participant in each of the three study groups. S35 on the other hand was the only environmental isolate that could not resist serum killing. Out of the three clinical strains used in the study, only one (C22) survived the bactericidal action of human serum from diabetic patients. From Table 5, the strains showed varying levels of susceptibility to bactericidal action of serum. After exposure to serum, the percent survival of E30 was recorded at 100% demonstrating higher resistance to bactericidal activity of serum. The other strains recorded percent survival rates ranging from 3-80%.

**Table 4:** Serum Survival of Environmental and Clinical Isolates

| Strain ID | Source | Serogroup | Diabetic patients (30) | Liver disease patients (10) | Healthy Controls (10) |
| --- | --- | --- | --- | --- | --- |
| S1 | Environmental | non-O1/O139 | - | 3 | - |
| S35 | Environmental | non-O1/O139 | - | - | - |
| E4 | Environmental | non-O1/O139 | 4 | 2 | 1 |
| E19 | Environmental | non-O1/O139 | 2 | 1 | - |
| E30 | Environmental | non-O1/O139 | - | 1 | - |
| E32 | Environmental | non-O1/O139 | - | 2 | - |
| C6 | Clinical | O1 | - | - | - |
| C17 | Clinical | O1 | - | - | - |
| C22 | Clinical | O1 | 2 | - | - |

Footnote: "numeric value" indicates frequency of survival in a specific study group. "-" indicates no survival.

**Table 5.** Rate of Serum Survival

| Survived strains/sample ID<sup>a</sup> | T<sub>0</sub> (Log CFU) | T<sub>4</sub> (Log CFU/ml ± standard deviation)<sup>b</sup> | % Survival of bacterial cells in serum after 4h incubation [(T<sub>4</sub>/T<sub>0</sub>) ×100] |
| --- | --- | --- | --- |
| E30/LD4 | 2.4×10<sup>6</sup> | 2.8×10<sup>6</sup> | 100% |
| E19/LD4 | 4.1×10<sup>6</sup> | 3.3×10<sup>6</sup> | 80% |
| E19/DB16, E19/DB19 | 4.1×10<sup>6</sup> | 2.6×10<sup>5</sup> – 1.1×10<sup>6</sup> | 6-26% |
| E4/DB4, E4/DB5, E4/DB8, E4/DB29 | 3.2×10<sup>6</sup> | 1.8×10<sup>5</sup> – 8.6×10<sup>5</sup> | 6-27% |
| E4/LD7, E4/LD8 | 3.2×10<sup>6</sup> | 4.6×10<sup>5</sup> - 1.3×10<sup>6</sup> | 14-41% |
| E4/HC5 | 3.2×10<sup>6</sup> | 3.2×10<sup>5</sup> | 10% |
| E32/LD7, E32/LD8 | 2.0×10<sup>6</sup> | 2.0×10<sup>5</sup> – 2.5×10<sup>6</sup> | 10-100% |
| S1/LD7, S1/LD8, S1/LD9 | 2.5×10<sup>6</sup> | 8.0×10<sup>4</sup> - 1.0×10<sup>6</sup> | 3-40% |
| C22/DB11, C22/DB15 | 2.4×10<sup>6</sup> | 8.4×10<sup>5</sup> - 1.0×10<sup>6</sup> | 35-42% |

Footnote: \*LD= Liver disease \*DB= Diabetic \*HC= Healthy Control.

**Table 6:** Virulence Genes Distribution among Clinical and Environmental Strains

| Strain ID | Source | Strain | Serogroup | Virulence Genes |  |  |  |  |  |  |  |  |  |  |  |  |  |  |  |
| --- | --- | --- | --- | --- | --- | --- | --- | --- | --- | --- | --- | --- | --- | --- | --- | --- | --- | --- | --- |
|  |  |  |  | *ctxA* | *ctxB* | *zot* | *ace* | *tcpA* | *hlyA* | *chxA* | *rtxA* | *toxR* | *als* | *mshA* | *T6SS* | *ompT* | *VgrG* | *entD* | *FepE* |
| C22 | Clinical | *V. cholerae* | O1 | + | + | + | + | + | + | - | + | + | + | + | + | + | + | - | - |
| C6 | Clinical | *V. cholerae* | O1 | + | + | + | - | + | + | - | + | + | + | + | + | + | + | - | - |
| C17 | Clinical | *V. cholerae* | O1 | + | + | + | - | - | + | - | + | + | + | + | + | + | + | - | - |
| S1 | Environmental | *V. cholerae* | non O1/O139 | - | - | - | - | - | + | - | + | + | + | + | - | - | - | + | - |
| S35 | Environmental | *V. cholerae* | non O1/O139 | - | - | - | - | - | + | - | + | + | + | - | + | - | - | - | - |
| E4 | Environmental | *V. cholerae* | non O1/O139 | - | - | - | - | - | + | + | + | + | + | - | - | - | - | - | + |
| E19 | Environmental | *V. cholerae* | non O1/O139 | - | - | - | - | - | + | - | + | + | + | - | - | - | - | + | - |
| E30 | Environmental | *V. cholerae* | non O1/O139 | - | - | - | - | - | + | - | - | + | + | + | - | - | - | + | - |
| E32 | Environmental | *V. cholerae* | non O1/O139 | - | - | - | - | - | + | - | + | + | + | + | - | - | - | + | - |

Footnote: “-” absence of genes, “+” presence of genes, “+” genes exclusively associated with environmental strains. Sequence data deposited in NCBI with the Accession numbers: SAMN26804331, C22; SAMN26804327, C6; SAMN26804329, C17; SAMN26804320, S1; SAMN26804321, S35; SAMN26804322, E4; SAMN26804323, E19; SAMN26804324, E30; SAMN26804325, E32.

To evaluate the contribution of the complement cascade in the lysis or resistance phenotype of *V. cholerae* strains, sera from selected individual volunteers were heat inactivated (56°C for 30 min) prior to bacterial exposure. The clinical strain, C22 survived and showed significant growth (2×10^4^ - 2.56×10^6^ CFU/mL) in inactivated serum of both diabetic (DB29) and liver (LD7) disease patients compared to no bacterial growth without heat inactivation of the native sera. In the case of the environmental isolates, E4 showed more than 100-fold increase in serum survival rate with DB29 and LD7 patients, including the serum of a healthy individual (HC2) following heat inactivation (data not shown).

The assigned numeric value to the sample ID refers to patient number. ^b^Values presented are log numbers of the means of bacterial growth after exposure to serum as measured by plate count (CFU/mL). Experiment for each strain was performed in three technical replicates.

### 3.3. The cholix toxin (*chxA*), genes encoding for siderophores (*fepE, EntD*) are potential virulence factors used by *V. cholerae* to survive outside the gastrointestinal tract and in human blood

The genomes of *V. cholerae* strains that survived bactericidal action of serum were analysed for genes that may contribute to their survival using the rapid annotation subsystem analysis (RASA) bioinformatic tool. Particular emphasis was placed on the virulence, iron acquisition and stress response gene contents of the pathogens. Table 5 shows the virulence genes distribution among the *V. cholerae* isolates after pangenome analysis. The clinical isolates possessed intact toxigenic genes (*ctxA, ctxB, tcpA*) whilst the environmental isolates lacked these cholera toxin genes required for pandemic cholera spread. However, the environmental strains contained unique genes that may play a significant role in *V. cholerae* bacteremia. The E4 strain was found to possess the cholix toxin (*chxA*) gene, which is known to be responsible for extraintestinal infections [23]. Over 83% of the environmental isolates possessed genes encoding for siderophores (*fepE*, and *entD*) normally used by bacterial pathogens to scavenge for iron in a limited environment for survival. Other genomic features of the strains can be found in our recent publication [24].

### 3.4. IL-1ß, IL-13 and RANTES are the immune cell components elicited during *V. cholerae* bacteremia

To evaluate the impact of *V. cholerae* on host immune response, 25 human cytokines (IL-10, IL-12, IL-13, IL-15, IL-17A, IL-1 beta, IL-IRA, IL-2, IL-2R, IL-4, IL-5, IL-6, IL-7, IL-8, IP-10, MCP-1, MIG, MIP-1 beta, MIP-1 alpha, RANTES, TNF alpha) were assayed. Only IL-1ß, IL-13 and RANTES were significantly expressed after 24 hours stimulation of PBMCs with *V. cholerae* (Fig 1).

**Fig 1.**
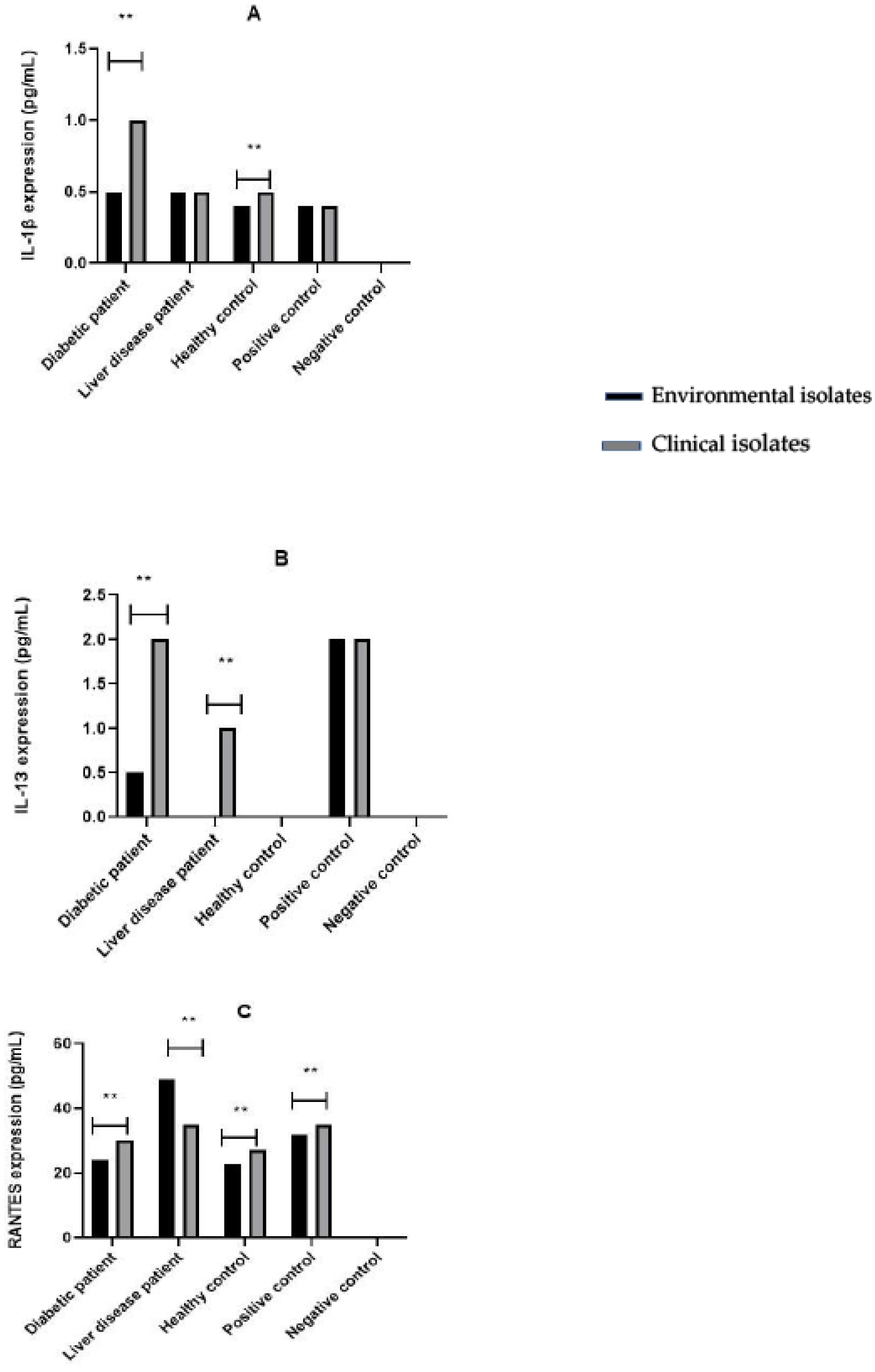
Cytokine Expression Levels After 24 hr Stimulation of PBMCS with *V. cholerae*. (A) IL-1ß expression by PBMCs after stimulation with environmental and clinical isolates. (B) IL-13 expression after stimulation with environmental and clinical isolates. (C) RANTES expression after stimulation with environmental and clinical isolates. [**] indicates variables with statistical differences using the GraphPad Prism statistical tool. Statistical significance was determined using the 2-way ANOVA test (P<0.05). Each bar represents the means ± SD from two independent experiments.

The clinical strains of *V. cholerae* were able to induce the production of higher levels of IL-1ß in diabetic patients (1pg/ml) compared to the environmental strains. However, there was no significant differences in expression levels between clinical and environmental strains in liver disease patients (0.5pg/ml) as well as the healthy controls. IL-13 expression was highly induced by clinical isolates in both diabetic (2pg/ml) and liver disease patients (1pg/ml) compared to environmental strains while there was no significant expression observed in healthy controls. Appreciable RANTES expression levels were also observed after stimulation by both environmental and clinical isolates in both diabetic and clinical patients.

## 4. Discussion

Cholera pathogens are generally considered to be non-invasive intestinal organisms linked with varied degrees of gastroenteritis [25]. The non-O1/non-O139 serogroups have been associated with extraintestinal infections such as septicemia, wound infection, skin infection, cholecystitis and meningitis despite being biochemically indistinguishable from toxigenic *V. cholerae* O1/O139 that causes cholera illness [26]. In recent years, the number of non-O1/non-O139 *Vibrio cholerae* bacteremia cases has been expanding across the globe [27-30]. Invasive infections induced by non-O1/non-O139 *V. cholerae* are rare yet deadly. Deshayes et al. [4] reported that one-third of patients died from non-O1/non-O139 *V. cholerae* bacteremia infections.

This study aimed to identify host and pathogen genetic factors contributing to *V. cholerae* survival outside the gastrointestinal tract by screening clinical and environmental strains of *V. cholerae* in human serum. Out of the 50 samples that were screened *in*-*vitro*, 36% of individuals were susceptible to *V. cholerae* bacteremia. These patients included individuals who were clearly diagnosed with hepatitis B cirrhosis and diabetes making them potentially prone to *V. cholerae* bacteremia as reported from other studies [7, 27]. The results indicate that non-O1/O139 strains of *V. cholerae* have the higher tendency of surviving in human sera compared to the toxigenic clinical strains carrying the *ctxA* and *ctxB* genes. Normal human serum is bactericidal for most types of bacteria because of its main component complement factors. The contribution of the complement cascade in regulating host susceptibility to bacterial infections is well documented [31]. As a result, resistance to serum killing is regarded as an essential virulence feature of invasive bacteria [32]. The observed increased survival rate of *V. cholerae* in serum following heat inactivation in this study suggests that complements play a role in the serum lysis process. Recent studies have proven that heat inactivated sera contained reduced amounts of complement factors [17], which mimics conditions in immunosuppressed patients with impaired innate and adaptive immune functions.

WGS analysis revealed the exclusive possession of the cholix toxin (*chxA*), enterobactin synthase component D (*entD*) and ferric enterobactin transport protein (*FepE*) genes by the environmental strains (Table 4). Studies by Awasthi et al. [23] suggested that the cholix toxin is responsible for extraintestinal infections in *V. cholerae*. Purdy et al. [33] and Jørgensen et al. [34] have previously shown that the environmental non-O1/O139 can carry a gene encoding a toxin that is similar in sequence and activity to that of the exotoxin A from *Pseudomonas aeruginosa*. The high structural similarity between exotoxin A and cholix toxin suggests that cholix toxin is likely to gain access to eukaryotic host cells via receptor-mediated endocytosis [23]. Purdy et al. [35] further posited that cholix toxin plays a major role in the fitness of *V. cholerae* which probably explains the hypervirulence traits of the strain (E4) that carried this toxin. Our genome analysis of survived strains also identified genes (*entD, FepE*) encoding for siderophores. It is well known that Iron (III) is an essential element for bacterial survival due to its involvement in many cellular processes [36]. Pathogenic bacteria have therefore developed strategies to scavenge and absorb irons from their surroundings including from host’s storage/transport. Several studies indicate that siderophores protect microorganisms from metal toxicity and thus may confer virulence-associated gains to pathogens during infections [37, 38].

Although several immune signal molecules including cytokines and chemokines may play an important role in disease pathogenesis, only IL-1ß, IL-13 and RANTES were detected at appreciable levels after stimulation of PBMCs with *V. cholerae* (Figure 3 A-C). IL-1ß is a proinflammatory cytokine produced by immune cells during inflammation or infection. Generally, IL-1ß acts to promote antigen-specific immune responses, inflammation and the remodelling of the extracellular matrix [39]. The data suggests that the clinical strains of *V. cholerae* are more capable of inducing IL-1ß production in diabetic patients which was not the case in patients suffering from liver diseases. On the other hand, the clinical strains induced IL-13 expression, an anti-inflammatory cytokine, in both diabetic and liver disease patients, consistent with findings from other studies [40]. The level of expression of IL-13 by environmental strains was however low in diabetic patients and absent in both liver disease patients and healthy controls. This difference in host responses may be mediated by the presence of the cholera toxin. *In vitro* studies have demonstrated the pre-treatment with cholera toxin suppresses the induction of pro-inflammotry cytokines in LPS-stimulated macrophages, suggesting that *V. cholerae* toxins have some anti-inflammatory activity [41]. The environmental strains perhaps have an unknown mechanism of evading cell mediated host immune responses and thus, their higher survivability in serum.

We observed higher level of expression of RANTES, a chemokine responsible for attracting many cell types including T cells, eosinophils and basophils to sites of infection. There was no appreciable variation in RANTES expression levels between PBMCs stimulated with environmental and clinical isolates. Studies have shown that in RANTES-stimulated T cells, there is an induction and up-regulation of specific adhesion and activation molecules in a lymphocyte function–associated antigen pathway. This suggests that once these cells are activated, their ability to respond to bacterial or viral infections may be significantly enhanced [42]. Other evidence suggests that RANTES reduces histamine release, halting the inflammatory response and redirecting the cellular surroundings toward wound repair and healing [43]. RANTES elevation has also been linked to infection severity, tissue damage, and mortality in sepsis [44], especially with viral infections. Further research is however needed to shed more light on the importance of RANTES in immunological responses to *V. cholerae* infection and to demonstrate its utility as a biomarker for monitoring severity of immunological reactions.

## 5. Conclusions

This research revealed an intriguing finding that could help better understand the evolution of *V. cholerae* and the molecular pathways involved in *V. cholerae* bacteremia. Although, we were limited in terms of the number of participants and bacterial isolates used, the study clearly demonstrates the chances of some non-O1/non-O139 *V. cholerae* strains causing bloodstream infections, particularly among immunocompromised individuals. The environmental strains used in this study were identified to possess the cholix toxin gene and genes encoding for siderophores (*fepE, entD*). The identification of these genes in the *V. cholerae* isolates suggest that their presence in blood could be prolonged by scavenging for iron from their host leading to severe infections. To explore further on this, it may be necessary to generate in-frame deletions in these genes and testing the mutant strains for serum resistance. Gene expression profiling of serum resistant *V. cholerae* strains and expressed cytokines will also help validate the genes responsible for *V. cholerae* bacteremia and the specific immune response elicited during the infection process. Finally, public health officials and clinicians should be aware of the risk of *V. cholerae* bacteremia because of the rare nature of the infection which could be missed in routine laboratory diagnosis leading to fatal outcomes.

## Acknowledgement

Analysis was performed using computing resources provided by University of Ghana and the West African Centre for Cell Biology of Infectious Pathogens (WACCBIP). We are thankful to Enoch Kofi Amoako (WACCBIP) for his assistance with the analysis of the whole genome sequence data. We are also thankful to Emmanuel Amoako Amposah for his helpful assistance in the laboratory analysis.

